# Autophagic flux is increased in peripheral blood mononuclear cells in atherosclerotic vascular disease and associates inversely with adverse cardiovascular events

**DOI:** 10.64898/2026.09.21.753342

**Authors:** Bronwyn Beelders, Leanne K. Hein, Naomi Wattchow, Mau T. Nguyen, Shabnam Torabiardakani, Thalia Salagaras, Sanuri Liyanage, Christina A. Bursill, Robert Fitridge, Neil McMillan, Christopher L. Delaney, Timothy J. Sargeant, Peter J. Psaltis

## Abstract

**Background:** Autophagy is a homeostatic pathway supporting stress adaptation and is dysregulated in atherosclerosis. Its potential as a biomarker or therapeutic target in atherosclerotic vascular disease (ASVD) remains incompletely defined. We measured autophagic flux in peripheral blood mononuclear cells (PBMCs) from patients with peripheral arterial disease (PAD) or carotid stenosis (CS), compared with healthy controls, and explored clinical outcome associations.

**Methods:** Ninety-four patients with PAD or CS and 19 healthy controls were studied. Autophagic flux was quantified from fresh blood using a validated *ex vivo* chloroquine-inhibition ELISA measuring LC3B-II accumulation. Major adverse cardiovascular events (MACE) and major adverse limb events (MALE) were ascertained over a median follow-up of 828 days.

**Results:** The ASVD cohort comprised claudication (n = 16), chronic limb-threatening ischemia (CLTI; n = 49), and CS (n = 29). Autophagic flux was higher in ASVD than controls (mean 281.4 vs. 182.3 ng LC3B-II/mg protein/h; p < 0.0001) and remained independently associated after multivariable adjustment. Within CLTI, concurrent infection was associated with lower flux (p = 0.001), approaching control levels (p = 0.327). In CLTI, higher flux quartiles were associated with lower MACE risk, most strongly for quartile 3 (hazard ratio 0.07 vs. quartile 1, 95% CI 0.01–0.50; p = 0.009).

**Conclusion:** Autophagic flux is elevated in PBMCs from ASVD patients, independent of age and sex. Attenuated flux in CLTI with concurrent infection may indicate autophagic exhaustion in advanced disease. The association between higher flux and lower MACE in CLTI suggests prognostic utility, warranting evaluation in larger prospective studies.

## BACKGROUND

Atherosclerosis is a systemic, chronic inflammatory disease that affects multiple vascular territories through shared pathological mechanisms and risk factors. Peripheral arterial disease (PAD) and carotid stenosis (CS) are two clinically important manifestations, together imposing a substantial burden on patients and healthcare systems. PAD can cause claudication, rest pain, tissue loss, limb loss, and impaired quality of life, while CS is a major modifiable contributor to ischemic stroke. Both conditions are closely associated with coronary artery disease and carry high rates of cardiovascular morbidity and mortality ^1, 2^. Despite this burden, circulating biomarkers that reflect disease activity or predict adverse outcomes remain limited, and therapeutic advances beyond traditional risk-factor modification and revascularization have been relatively modest.

Autophagy is a highly conserved intracellular degradation pathway that maintains cellular homeostasis by targeting damaged organelles, misfolded proteins, and redundant cytoplasmic constituents for lysosomal degradation and recycling ^3^. Basal autophagic activity occurs in virtually all mammalian tissues and is upregulated by diverse cellular stressors, including oxidative stress, inflammation, hypoxia, and lipid excess, all of which are prominent within atherosclerotic plaques ^4^. Through this adaptive response, autophagy supports cell survival, regulates lipid metabolism, modulates immune cell phenotype, preserves efferocytosis, and attenuates inflammatory signaling ^5–9^. When autophagy is impaired, these protective mechanisms may be disrupted, promoting atherosclerotic progression and plaque destabilization. For example, impaired macrophage autophagy reduces efferocytosis and increases susceptibility to apoptosis, contributing to necrotic core expansion and sustained plaque inflammation ^5, 10^. Similarly, impaired autophagy in vascular smooth muscle cells increases plaque growth and instability with intraplaque hemorrhage in preclinical models ^11, 12^.

The biological effect of autophagy in any given context depends on the balance between autophagy induction and downstream degradative capacity ^13^. Only a small number of studies have examined autophagy in circulating immune cells in atherosclerotic vascular disease (ASVD), and most have reported reductions in autophagy-related gene expression or protein markers in PAD and coronary artery disease ^14–16^. However, these approaches rely on static measurements that cannot distinguish impaired autophagosome formation from impaired lysosomal degradation and therefore do not capture autophagic flux under physiologically relevant conditions. In addition, transcript abundance of autophagy-related genes does not necessarily correlate with functional autophagy ^17^. Building on a validated *ex vivo* method that preserves physiological autophagic activity in freshly isolated peripheral blood mononuclear cells (PBMCs) ^18^, this study aimed to compare autophagic flux in circulating immune cells from patients with ASVD and healthy controls. We also explored the relationship between autophagic flux and subsequent adverse cardiovascular and limb outcomes.

## METHODS

### Study design and population

This prospective cohort study recruited ASVD patients with a confirmed diagnosis of PAD or CS managed in an inpatient or outpatient setting at the Royal Adelaide Hospital or Flinders Medical Centre in Adelaide, Australia, under ethics approval (HREC/18/CAHLN/95, R20180206; 2023/HREC00203). Healthy control participants were recruited from Jones Radiology at the South Australian Health and Medical Research Institute under a different ethics application (HREC17115; ACTRN12623000458639). Controls were eligible if they had undergone coronary computed tomography angiography showing no coronary atherosclerosis and reported no history of PAD or CS on a standardized health questionnaire. All participants provided written informed consent prior to enrolment.

### PBMC isolation and ex vivo autophagic flux assay

Peripheral venous blood (10 mL) was collected into lithium heparin tubes and processed within 30 minutes of collection. Whole blood was divided equally into two tubes and incubated with 150 μM chloroquine (lysosomal inhibitor) or vehicle for one hour at 37°C with rotation.

Following incubation, blood was diluted 1:1 with cold phosphate-buffered saline (PBS). PBMCs were isolated by density-gradient centrifugation over Lymphoprep (Serumwerk Bernburg, Bernburg, Germany), followed by sequential red blood cell lysis, PBS washing, and pelleting by centrifugation. Paired PBMC pellets (treated and untreated) were snap-frozen on dry ice and stored at −80°C until analysis ^19^.

After thawing, PBMC pellets were resuspended in PBS containing 0.05% saponin to remove cytosolic microtubule-associated protein 1 light chain 3 beta-1 (LC3B-I), washed again and then resuspended in cell extraction buffer (Cell Signalling Technology, #35172) containing protease inhibitors. Samples were sonicated twice on ice for 20 seconds before being clarified.

Autophagic flux was quantified as the increment in LC3B-II concentration (ng/mg of protein) between chloroquine-treated and vehicle-treated samples, measured by commercial ELISA (Cell Signalling Technology, #35172), with the addition of a standard curve (0-5 ng/well) using recombinant human LC3B (Abcam, ab103506) to give a final unit of ng of LC3B-II/mg protein/h. This approach has been previously validated as a method for quantifying autophagic flux under physiological conditions in freshly isolated human PBMCs ^19^.

### Clinical outcomes

A retrospective review of electronic medical records was performed in July 2025 to ascertain incident clinical events from enrolment to the date of review. Major adverse cardiovascular events (MACE) were defined as a composite of non-fatal myocardial infarction, non-fatal stroke, and all-cause mortality. Major adverse limb events (MALE) were defined as a composite of acute limb ischemia, major amputation, or surgical/endovascular re-intervention for progressive limb ischemia. Outcomes were modelled as time to first event; recurrent events of the same type within an individual were not captured or analyzed separately.

### Statistical analysis

This was a pilot exploratory study. The control sample size was estimated using the mean basal autophagic flux reported by Bensalem *et al.* (299.3 ± 95.4 ng LC3B-II/mg protein/hour) ^20^. Assuming a minimum clinically relevant difference of 30% in the ASVD group, this produced a suggested sample size of 15 controls, providing 90.7% power at α = 0.05 at the study’s planned allocation ratio of approximately five ASVD patients per control.

Continuous variables are presented as mean (95% confidence interval) or median [interquartile range (IQR)] according to distributional assessment by the Shapiro-Wilk test, and categorical variables as counts and proportions. Between-group comparisons of continuous variables were performed using the independent-samples t-test or one-way analysis of variance (ANOVA) for normally distributed data, and the Mann-Whitney U test or Kruskal-Wallis test for non-normally distributed data. Categorical variables were compared using the χ² test or Fisher exact test as appropriate.

The independent association between autophagic flux and patient-level variables was examined by univariable and multivariable linear regression. Autophagic flux was log-transformed prior to regression to satisfy the normality assumption of residuals. The multivariable model was prespecified to include age, sex, comorbidities [diabetes mellitus, hypertension, hypercholesterolemia, chronic kidney disease (CKD, stage ≥3; eGFR <60 mL/min/1.73 m²), body mass index (BMI), active smoker, and medication use (statin intensity). Associations between autophagic flux and time-to-event outcomes (MACE and MALE) were assessed using Cox proportional hazards regression, with flux modelled as quartiles (Q1, lowest, as reference category). All Cox models included age, sex, diabetes mellitus, hypertension, current smoking, and concurrent infection as prespecified covariates. All hypothesis tests were two-sided with a significance threshold of α = 0.05. Figures were generated and statistical analyses were performed using GraphPad Prism (Version 11.0.2, GraphPad Software, Boston, Massachusetts USA), IBM SPSS Statistics (Version 31.0.2.0, IBM Corp, Armonk, New York USA) and Stata Statistical Software (Version 19.5 Stata Corp., College Station, Texas USA). Given the small number of events, all outcome analyses should be considered exploratory.

## RESULTS

### Participant characteristics

113 participants were enrolled: 94 with ASVD and 19 healthy controls. Within the ASVD group, 65 patients had PAD [49 chronic limb threatening ischemia (CLTI), 16 claudication] and 29 had CS. The ASVD group was significantly older than the control group (mean 71 vs. 53 years; p < 0.001) and included a higher proportion of males (76.6% vs. 47.4%; p = 0.010). Diabetes mellitus was also more prevalent in the ASVD group (**Table 1**). A breakdown of characteristics by ASVD subgroup is provided in **Supplementary Table 1**.

**Table 1.** Baseline participant characteristics.

|  |  | **Control**<br>**(n=19)** | **ASVD**<br>**(n=94)** | **Difference**<br>**(p-value)** |
| --- | --- | --- | --- | --- |
| **Demographics** |  |  |  |  |
| Age, years, mean (SD) |  | 53 (8.6) | 71 (9.0) | <0.001 |
| Male sex, n (%) |  | 9 (47.4) | 72 (76.6) | 0.010 |
| **Comorbidities** |  |  |  |  |
| Diabetes mellitus, n (%) |  | 2 (10.5) | 47 (50.0) | 0.002 |
| Hypercholesterolemia, n (%) |  | 11 (62.5) | 48 (51.1) | 0.59 |
| Hypertension, n (%) |  | 8 (42.1) | 48 (51.1) | 0.48 |
| CKD (≥ Stage 3), n (%) | | 0 (0) | 5 (5.3) | 0.30 |
| Active smoker, n (%) |  | 3 (15.8) | 25 (26.6) | 0.32 |
| **Medication** |  |  |  |  |
| Statin, n (%) * | Nil | 12 (63.2) | 14 (14.9) | 0.36 |
|  | Low | 0 (0) | 0 (0) |  |
|  | Moderate | 4 (21.1) | 17 (18.1) | 0.39 |
|  | High | 3 (15.8) | 63 (67.0) | 0.47 |
| Metformin, n (%) |  | 2 (10.5) | 35 (37.2) | 0.024 |

Abbreviations: ASVD: atherosclerotic vascular disease; CKD: chronic kidney disease (stage $\geq 3$ ; eGFR $< 60$ mL/min/1.73 m<sup>2</sup>); SD: standard deviation. \*Low-intensity statin use defined as simvastatin 10 mg or pravastatin 10-20 mg; moderate-intensity as atorvastatin 10-20 mg, rosuvastatin 5-10 mg, simvastatin 20-40 mg or pravastatin 40-80 mg; high-intensity as atorvastatin 40-80mg, rosuvastatin 20-40mg

### Basal LC3B-II abundance and autophagic flux

LC3B is a ubiquitin-like protein that participates in autophagosome biogenesis and autophagic cargo selection; LC3B-II is the lipidated, membrane-bound form that marks autophagosomes^21^. Basal LC3B-II concentration did not differ significantly between the ASVD group (median 184.2 ng LC3B-II/mg protein, IQR 124.2–234.4) and healthy controls (median 156.1 ng LC3B-II/mg protein, IQR 141.3–206.8; p = 0.70), indicating comparable steady-state autophagosome abundance between groups (**Figure 1A**).

**Figure 1.**
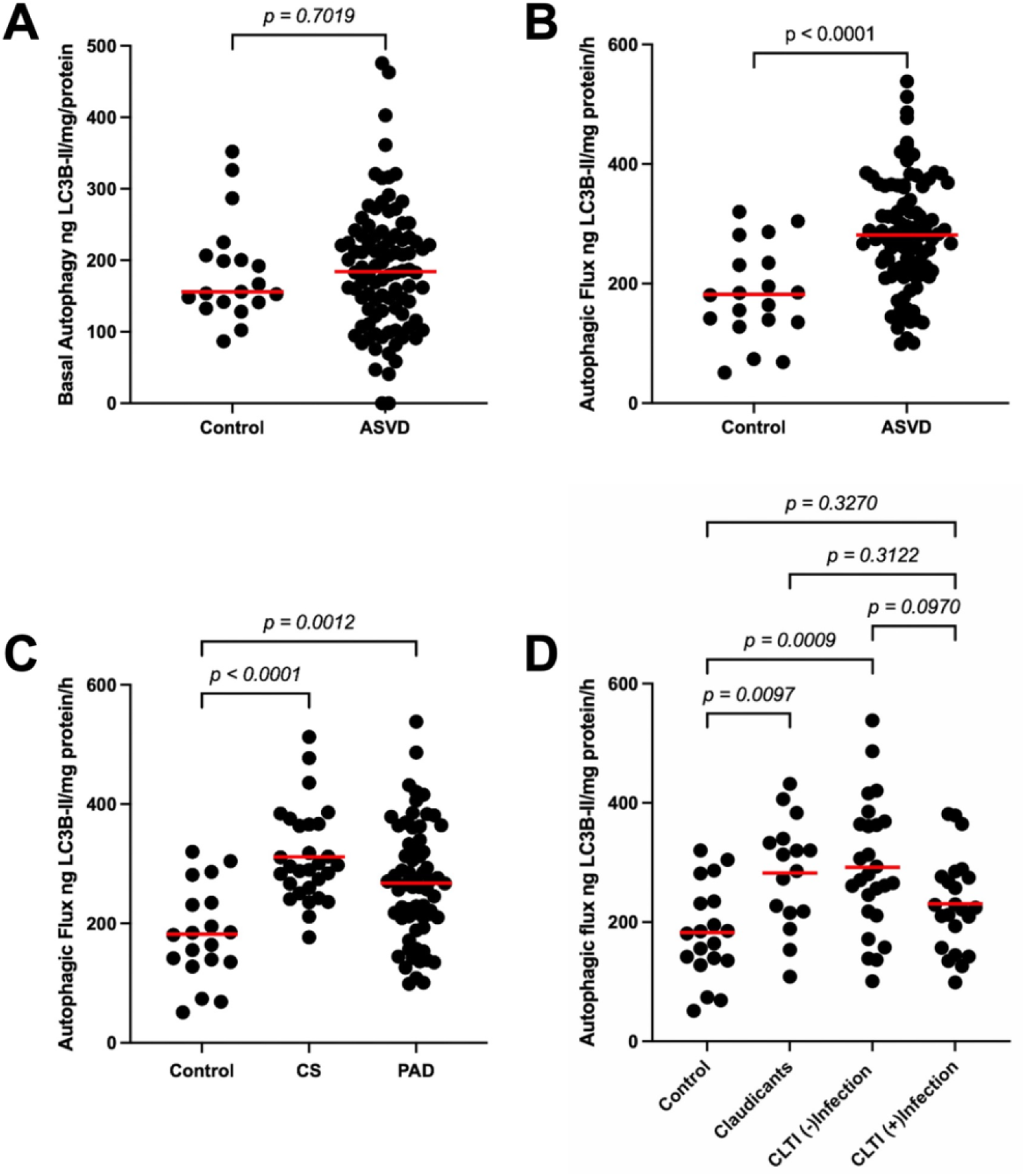
Basal LC3B-II concentration and autophagic flux in peripheral blood mononuclear cells (PBMCs) across atherosclerotic vascular disease (ASVD) subgroups. (A) Basal LC3B-II concentration in PBMCs from healthy controls (n = 19) and patients with ASVD (n = 94). (B) Autophagic flux in healthy controls compared with patients with ASVD. (C) Autophagic flux in healthy controls, patients with peripheral artery disease (PAD; n = 65) and patients with carotid stenosis (CS; n = 29). (**D**) Autophagic flux in healthy controls, patients with claudication (n = 16), patients with chronic limb-threatening ischemia (CLTI) without soft-tissue infection (n = 25), and patients with CLTI complicated by infection (n = 24). Red bars indicate the group median (A) or mean (B–D). Statistical comparisons were performed using the Mann-Whitney U test (A), Welch’s t-test (B), and one-way ANOVA (C, D).

In contrast, autophagic flux - quantified as the chloroquine-induced increment in LC3B-II - was significantly higher in the ASVD group (mean 281.4 ng LC3B-II/mg protein/h, 95% CI 262.2-300.6) than in controls (mean 182.3 ng LC3B-II/mg protein/h, 95% CI 144.3-220.2; p < 0.0001) (**Figure 1B**). On subgroup analysis, flux did not differ significantly between patients with PAD and those with CS (PAD: mean 267.9 ng LC3B-II/mg protein/h, 95% CI 243.7-292.0; CS: mean 311.8 ng LC3B-II/mg protein/h, 95% CI 282.2–341.4; p = 0.077), and both subgroups had higher flux than controls (p = 0.0012 and p < 0.0001, respectively) (**Figure 1C**).

Among the 49 patients with CLTI, 24 had concurrent lower limb infections at the time of blood collection, as determined by clinical assessment and antibiotic treatment, while 25 did not. Further stratification of autophagic flux results within the wider PAD cohort demonstrated that both the claudication (mean 282.3 ng LC3B-II/mg protein/h, 95% CI 233.7–330.9; p = 0.0097) and CLTI without infection groups (mean 291.9 ng LC3B-II/mg protein/h, 95% CI 248.0–335.9; p = 0.0009) had higher flux values than controls. By contrast, CLTI patients with infection had flux levels that were not significantly different from controls (mean 230.6 ng LC3B-II/mg protein/h, 95% CI 196.6-264.6; p = 0.33) (**Figure 1D**).

### Multivariable determinants of autophagic flux

Univariable analysis demonstrated significant positive associations between autophagic flux and CS (p < 0.001), age (p = 0.036), male sex (p = 0.022), and statin use at moderate (p = 0.037) and high intensity (p < 0.001). After multivariable adjustment for age, sex, comorbidities and medication use, the presence of ASVD was the only variable that remained independently associated with autophagic flux. Each ASVD group demonstrated significantly higher adjusted flux relative to controls: CS showed the strongest association (β = 0.59, 95% CI 0.26-0.91; p = 0.001), followed by claudication (β = 0.50, 95% CI 0.19-0.83; p = 0.003) and CLTI (β = 0.35, 95% CI 0.02-0.68; p = 0.039). Age and sex were no longer independently associated with flux after adjustment (age: β = −0.003, 95% CI −0.01 to 0.01; p = 0.53; sex: β = −0.07, 95% CI −0.25 to 0.01; p = 0.48) (**Table 2**).

**Table 2.** Univariable and multivariable linear regression of log-transformed autophagic flux.

| Variable | Unadjusted $\beta$<br>(95%CI) | p-value | Adjusted $\beta$<br>(95%CI) | p-value |
| --- | --- | --- | --- | --- |
| **Disease Group†** |  |  |  |  |
| PAD |  |  |  |  |
| Claudicant | 0.55 (0.28-0.81) | <0.001 | 0.50 (0.19-0.83) | 0.003 |
| CLTI | 0.40 (0.18-0.61) | <0.001 | 0.35 (0.02-0.68) | 0.039 |
| Carotid Stenosis | 0.62 (0.38-0.85) | <0.001 | 0.59 (0.26-0.91) | 0.001 |
| **Demographics** |  |  |  |  |
| Age | 0.01 (0.00-0.02) | 0.036 | -0.00 ( -0.01–0.01) | 0.53 |
| Sex (male) | -0.21 (-0.39-0.03) | 0.022 | -0.07 (-0.25-0.01) | 0.48 |
| **Comorbidities** |  |  |  |  |
| Diabetes mellitus | 0.07 (-0.10-0.23) | 0.43 | -0.02 (-0.31-0.26) | 0.88 |
| Hypercholesteremia | 0.13 (-0.04-0.29) | 0.13 | 0.32 (-0.03-0.67) | 0.069 |
| Hypertension | 0.11 (-0.06-0.27) | 0.21 | -0.28(-0.63–0.07) | 0.12 |
| CKD (≥ Stage 3) | 0.25 (-0.15-0.65) | 0.22 | 0.37 (-0.03-0.76) | 0.070 |
| BMI | -0.00 (-0.02-0.01) | 0.65 | -0.00 (-0.01-0.01) | 0.90 |
| Active smoker | 0.09 (-0.10-0.28) | 0.35 | 0.05(0.13-0.24) | 0.57 |
| **Medication** |  |  |  |  |
| Metformin | 0.07 (-0.11-0.25) | 0.43 | -0.01 (-0.29-0.27) | 0.94 |
| Statin* |  |  |  |  |
| Moderate | 0.26 (0.02-0.50) | 0.037 | 0.17 (-0.09-0.43) | 0.20 |
| High | 0.37 (0.19-0.56) | <0.001 | 0.21 (-0.01-0.43) | 0.065 |

Autophagic flux was log-transformed. †Healthy control group as reference. \*Nil statin use as reference. Abbreviations: BMI: body mass index; CI: confidence interval; CKD: chronic kidney disease (stage $\geq 3$ ; eGFR $< 60$ mL/min/1.73 m<sup>2</sup>); CLTI: chronic limb-threatening ischemia; PAD: peripheral arterial disease.

### Autophagic flux and adverse clinical outcomes

Over a median follow-up of 828 days (IQR,788-876), the ASVD group had 17 occurrences of MACE and 24 of MALE in 36 patients. As shown in **Table 3**, most adverse events were concentrated in the CLTI subgroup, consistent with the more advanced systemic atherosclerotic burden and well-established poorer prognosis of these patients. The limited number of events in the claudication and CS subgroups precluded stable multivariable estimation. Therefore, we restricted outcome analyses to the CLTI cohort.

**Table 3.**
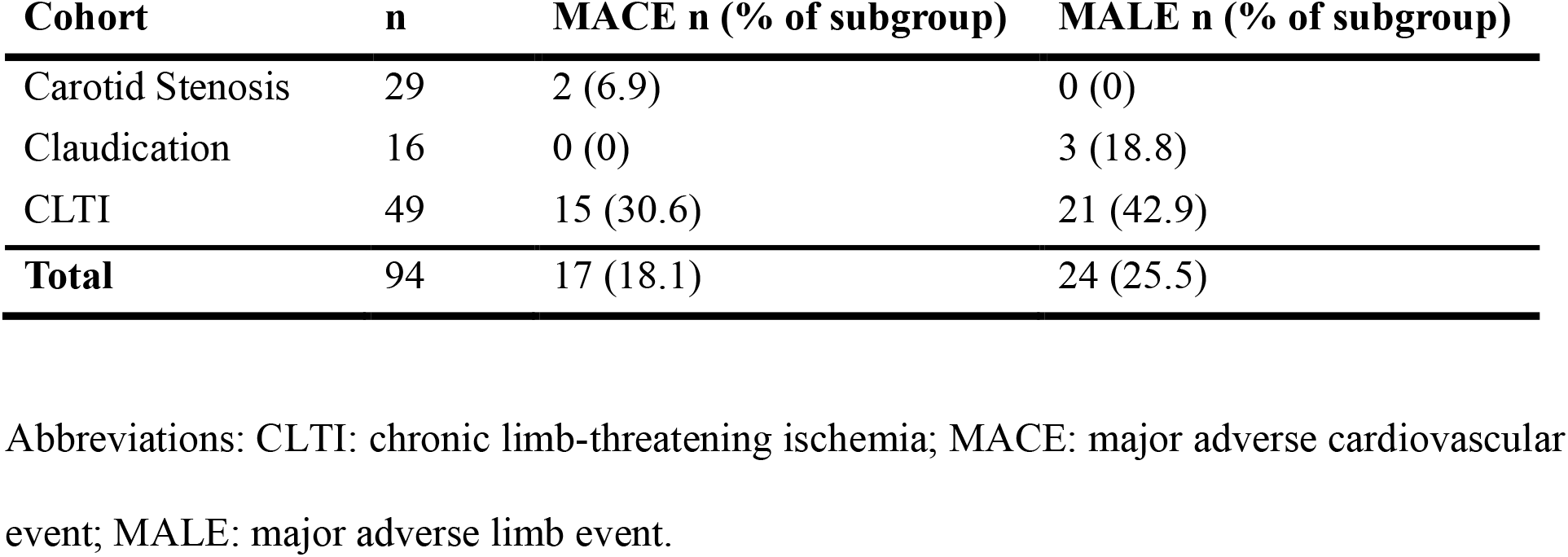
Incidence of adverse clinical events stratified by disease subgroup.

| **Cohort** | **n** | **MACE n (% of subgroup)** | **MALE n (% of subgroup)** |
| --- | --- | --- | --- |
| Carotid Stenosis | 29 | 2 (6.9) | 0 (0) |
| Claudication | 16 | 0 (0) | 3 (18.8) |
| CLTI | 49 | 15 (30.6) | 21 (42.9) |
| **Total** | **94** | **17 (18.1)** | **24 (25.5)** |

Abbreviations: CLTI: chronic limb-threatening ischemia; MACE: major adverse cardiovascular event; MALE: major adverse limb event.

In multivariable Cox proportional hazards regression, autophagic flux was modelled as quartiles, with quartile 1 (Q1) representing the lowest flux results and Q4 the highest. Higher flux quartiles were associated with significantly lower MACE hazard compared with Q1: Q2 HR 0.12 (95% CI 0.02–0.85; p = 0.034); Q3 HR 0.07 (95% CI 0.01–0.50; p = 0.009); and Q4 HR 0.18 (95% CI 0.04–0.81; p = 0.026) (**Figure 2A**). Diabetes mellitus was independently associated with higher MACE risk (HR 26.05, 95% CI 3.21–211.12; p = 0.002). Age, sex, smoking, and hypertension were not associated with MACE (data not shown).

**Figure 2.**
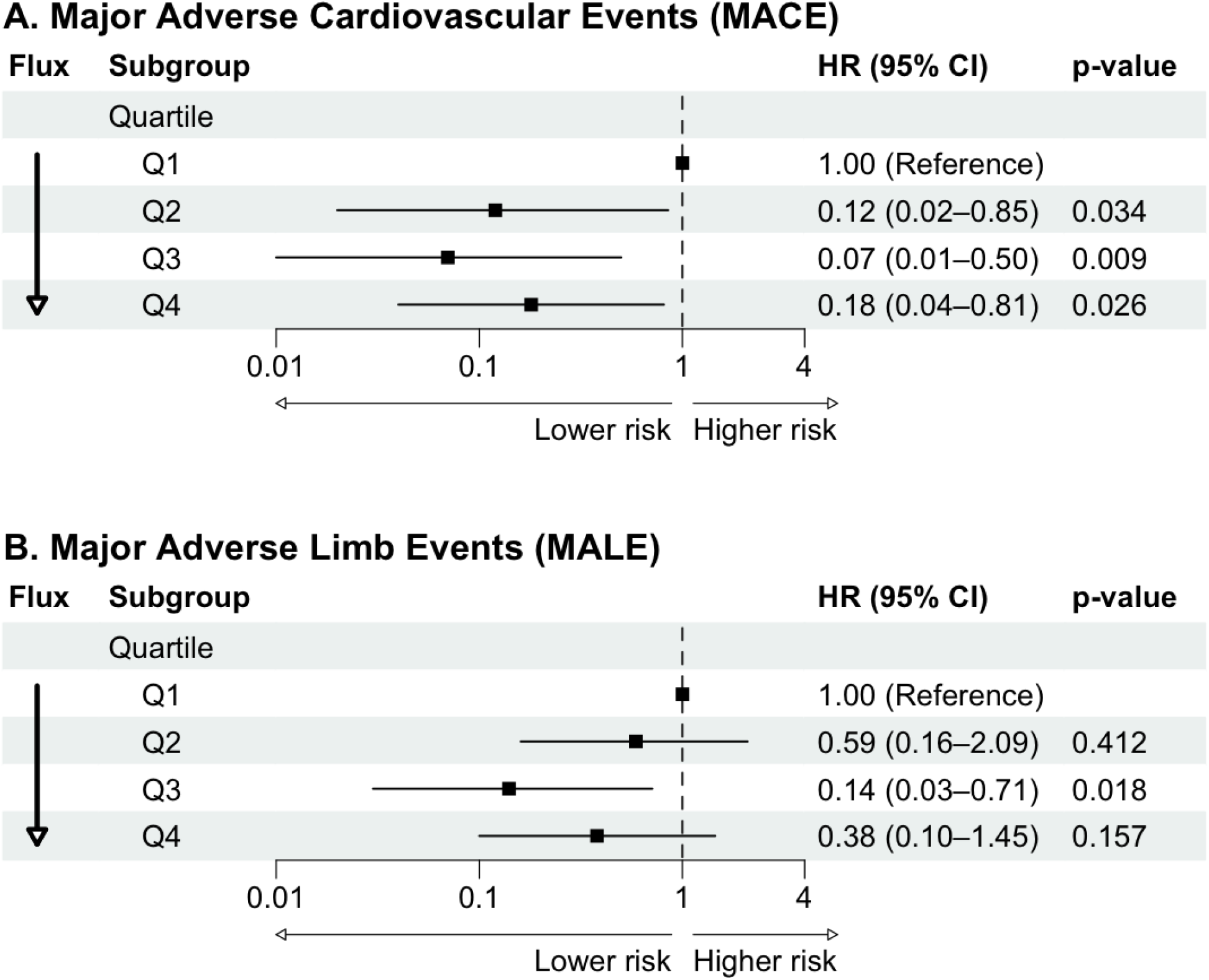
Forest plot of major adverse cardiovascular events (MACE) and major adverse limb events (MALE) by autophagic flux quartile. Adjusted hazard ratios (HR) with 95% confidence intervals (CI) are shown for **(A)** MACE and **(B)** MALE across quartiles of autophagic flux, with the lowest quartile (Q1) as the reference. Estimates were derived from multivariable Cox proportional hazards models adjusted for age, sex, diabetes mellitus, active smoking, hypertension, and concurrent infection. The vertical line denotes an HR of 1.0; values to the left indicate lower risk relative to Q1. CI, confidence interval; HR, hazard ratio; MACE, major adverse cardiovascular events; MALE, major adverse limb events; Q, quartile.

A similar but attenuated pattern was observed for MALE. The third quartile of autophagic flux was significantly protective compared to Q1 (HR 0.14, 95% CI 0.03–0.71; p = 0.018), while results for Q2 and Q4 were directionally consistent but did not reach statistical significance (Q2 vs Q1 HR 0.59, 95% CI 0.16–2.09; p = 0.41; Q4 vs Q1 HR 0.38, 95% CI 0.10–1.45; p = 0.16) (**Figure 2B**). None of the clinical covariates were independently associated with MALE.

## DISCUSSION

To our knowledge, this is the first clinical study to measure autophagic flux, rather than static autophagy markers, in PBMCs from patients with ASVD using a method validated to preserve physiological autophagic activity. Patients with ASVD, comprising PAD or CS, had higher autophagic flux than healthy controls, independent of age, sex, comorbid risk factors, and medication use. By contrast, basal LC3B-II abundance did not differ between groups, underscoring the added value of dynamic flux measurement over static protein quantification. Within the PAD cohort, autophagic flux showed progressive attenuation with increasing disease complexity, with the most advanced subgroup - CLTI with concurrent infection - demonstrating flux values that were no longer statistically different from controls. Notably, among patients with CLTI, higher autophagic flux quartiles were associated with a lower hazard of MACE, with a similar trend observed for MALE.

The existing clinical literature on autophagy in circulating immune cells in ASVD is limited and has largely described reductions in autophagy-related gene expression and protein markers. Dai *et al.* identified 20 autophagy-related genes that were downregulated in PBMCs from patients with PAD, together with reduced beclin-1 and LC3B-II protein levels ^14^. Similarly, in coronary artery disease, Wu *et al.* reported lower LC3B gene expression in circulating leucocytes compared with controls, a finding that remained strongly associated with disease after adjustment for diabetes and hypertension (OR 7.81, 95% CI 2.32-26.32; p = 0.01) and was accompanied by reduced LC3B-II protein abundance ^16^. These findings are also broadly consistent with the report by Zhao *et al.* of lower beclin-1 and LC3B expression in peripheral monocytes from patients with acute coronary syndrome compared with controls or patients with stable angina ^15^. Taken together, these studies suggest reduced autophagy-related marker expression in circulating immune cells in ASVD, which appears at first to contrast with the elevated flux observed in the present study.

This apparent discrepancy most likely reflects fundamental methodological differences. LC3B-II is widely used as a surrogate marker of autophagosome abundance, but it provides only a static snapshot of a dynamic process. Its abundance may increase because of enhanced autophagosome biogenesis or impaired lysosomal degradation; therefore, when interpreted in isolation, LC3B-II cannot reliably indicate whether overall autophagy pathway activity is increased or decreased ^13,22^. Static protein markers of autophagy and transcript abundance of autophagy genes are also known not to correlate consistently with functional autophagy ^13, 17^. In addition, transferring cells to culture medium before measurement substantially reduces autophagic activity, potentially introducing systematic downward bias in studies using cultured or cryopreserved cells ^23^. The present study used a validated *ex vivo* chloroquine-inhibition assay in whole blood, followed by PBMC isolation, to assess autophagic flux under physiological conditions. In this context, the elevated flux observed here is not necessarily inconsistent with prior studies; rather, it likely reflects a compensatory autophagic response to the systemic oxidative and inflammatory stress present in ASVD.

PBMCs are accessible surrogates for investigating systemic cellular responses in chronic disease, and their autophagy status has been associated with disease severity across several conditions. In chronic obstructive pulmonary disease, PBMC autophagy markers correlate inversely with lung function and positively with circulating IL-6, IL-8, and TNF-α ^24^. In Kawasaki disease, elevated autophagic flux and increased autophagosome counts have been observed in PBMCs; co-culture experiments further suggest that PBMCs can directly induce autophagy and inflammatory cytokine expression in coronary artery endothelial cells ^25^. These observations support the use of PBMCs as a minimally invasive and clinically tractable window into systemic autophagic activity.

The elevation of autophagic flux in the ASVD cohort is particularly noteworthy because these patients were substantially older than controls, and autophagic activity is generally considered to decline with advancing age ^4^. In this study, age was not an independent predictor of flux after multivariable adjustment, whereas disease group remained independently associated with flux. This suggests that ASVD-specific pathophysiological stimuli, rather than age alone, were driving the autophagic response measured here. The metabolic milieu of ASVD is characterized by elevated oxidative stress, chronic systemic inflammation, and dyslipidemia, all of which are recognized inducers of autophagy ^26–29^. Reactive oxygen species can directly activate autophagy as a protective mechanism to clear damaged mitochondria and limit propagation of oxidative injury ^30, 31^. PBMCs from patients with diabetes and dyslipidemia - both prevalent comorbidities in the ASVD cohort - also exhibit elevated reactive oxygen species and increased autophagic activity ^32^. Inflammatory mediators may further amplify this response through interconnected pathways. Cytokines regulate autophagy through multiple signaling mechanisms, while damage-associated molecular patterns, including high mobility group box 1, can directly stimulate autophagic activity. Circulating high mobility group box 1 levels are elevated in both PAD and CS and may also contribute to plaque progression ^33–36^. Together, these observations support a model in which sustained systemic metabolic and inflammatory stress drives compensatory upregulation of autophagic flux in circulating immune cells from patients with ASVD.

Within the PAD cohort, subgroup analysis revealed a gradient of autophagic activity that tracked disease severity. Claudication and CLTI without infection were both associated with significantly elevated flux relative to controls, whereas flux in patients with CLTI and concurrent infection was closer to control values. This latter finding is unlikely to represent restoration of homeostatic autophagic function, but more plausibly reflects autophagic exhaustion in the setting of advanced, multifactorial cellular stress. Chronic oxidative stress can compromise lysosomal membrane integrity and disrupt autophagosome-lysosome fusion, producing a state in which autophagosomes accumulate but cannot be efficiently resolved, and net flux falls despite ongoing upstream activation ^37, 38^. Infection may compound this process: bacterial products can directly interfere with autophagosome maturation and lysosomal fusion, while also inducing mitochondrial dysfunction that further limits autophagic capacity ^39–41^. The pattern observed here - elevated flux in early-to-intermediate disease, followed by attenuation in the most advanced infected subgroup - is therefore biologically consistent with a transition from adaptive autophagy to autophagic failure under sustained and compounding cellular stress.

The non-linear relationship between autophagic flux and MACE risk also merits consideration. Intermediate flux quartiles, particularly Q3, showed the most pronounced association with reduced risk, whereas the protective association was somewhat attenuated in Q4. Although event numbers were small and these analyses should be considered exploratory, this pattern is biologically plausible given the dual and context-dependent role of autophagy in atherosclerosis. Insufficient flux may allow cellular damage to accumulate and impair immune function, whereas very high flux may reflect a degree of systemic stress that independently confers risk.

Several limitations in our study warrant acknowledgement. First, the relatively small sample size, particularly within disease subgroups, restricted the statistical power of secondary analyses, and the outcome analyses are hypothesis-generating at best. Second, the control group was not age- or sex-matched to the ASVD cohort, reflecting the known epidemiology of atherosclerosis favoring older male individuals ^42, 43^. However, multivariable adjustment for both variables did not alter the independent association between disease group and autophagic flux. Third, autophagic flux was measured at a single time point; longitudinal sampling will be required to determine whether flux changes with disease progression, clinical events, or therapeutic intervention. Finally, the absence of concurrent tissue or imaging data precludes direct mechanistic correlation between systemic autophagic activity and local plaque characteristics.

## CONCLUSION

Patients with ASVD demonstrated elevated autophagic flux in circulating PBMCs, independent of age and sex. This increase was attenuated in patients with CLTI and concurrent infection, suggesting possible autophagic exhaustion in advanced disease. Within the CLTI cohort, higher autophagic flux was independently associated with a lower hazard of MACE. These findings suggest that PBMC autophagic flux, quantified from a small fresh blood sample using a validated *ex vivo* assay, may be a feasible biomarker of systemic atherosclerotic disease activity. Larger prospective studies incorporating longitudinal sampling, tissue correlation, and broader vascular disease subgroups are needed to validate and extend these findings.

## SUPPLEMNTARY MATERIALS

There is a supplementary table that provides participant characteristics by disease state Supplementary Table 1: Participant characteristics by disease state

## AUTHOR CONTRIBUTIONS

*Conceptualization*, T.J.S and P.J.P; *Methodology*, T.J.S and L.K.H.; *Formal Analysis,* B.B and M.T.N.; *Investigation*, S.T, B.B, L.K.H, T.S, and S.L; *Resources*, P.J.P and T.J.S; *Data Curation,* T.S and S.L; *Writing – Original Draft Preparation*, B.B; *Writing – Review & Editing*, L.K.H, N.W, S.T, N.M, C.L.D, R.F, C.A.B, T.J.S, and P.J.P; *Supervision*, C.L.D, T.J.S, P.J.P, R.F; *Funding Acquisition*, P.J.P

## INSTITUTIONAL REVIEW BOARD STATEMENT

This study was conducted in accordance with the Declaration of Helsinki and approved by the Southern Adelaide and Central Adelaide Clinical Human Research Ethics Committee, South Australia (Approval numbers: HREC/18/CAHLN/95, R20180206; 2023/HREC00203, HREC17115; ACTRN12623000458639)

## INFORMED CONSENT STATEMENT

Written informed consent for inclusion in this research was obtained from the patients.

## DATA AVAILABILITY

The raw data supporting the conclusions of this article will be made available by the authors on request

## ACKNOWLEDGEMENTS

We gratefully acknowledge the study participants.

