## Supplemental Table 1 for "Autophagic flux is increased in peripheral blood mononuclear cells in atherosclerotic vascular disease and associates inversely with adverse cardiovascular events"

**Supplementary Table 1. Participant characteristics by disease state.**

|  | | |  | | **Control (n=19**) | **Claudicant**  **(n=16)** | **CLTI**  **(n=49)** | **Carotid Stenosis**  **(n=29)** |
| --- | --- | --- | --- | --- | --- | --- | --- | --- |
| **Demographics** | | |  | |  |  |  |  |
| Age, Years, Mean (SD) | | | | | 53.6 (8.6) | 67 (10.6) | 72.1 (9.5) | 71.2 (6.8) |
| Sex, n (%) | | | Male | | 9 (47.4) | 12 (75.0) | 39 (79.6) | 21 (72.4) |
| **Comorbidities** | | |  | |  |  |  |  |
| Diabetes mellitus, n (%) | | | |  | 2 (10.5) | 4 (25.0) | 34 (69.4) | 9 (31.0) |
| Hypercholesterolemia, n (%) | | |  | | 11 (62.5) | 10 (62.5) | 22 (44.9) | 16 (55.2) |
| Hypertension, n (%) | | |  | | 8 (42.11) | 10 (62.5) | 22 (44.9) | 16 (55.2) |
| CKD (≥ Stage 3), n (%) | | |  | | 0 (0) | 0 (0) | 5 (10.2) | 0 (0) |
| Active smoker, n (%) | | |  | | 3 (15.8) | 5 (31.3) | 10 (20.4) | 10 (34.5) |
| **Medication** |  | | | |  |  |  |  |
| Statin, n (%) * | | Nil | | | 12 (63.2) | 2 (12.5) | 7 (14.3) | 5 (17.2) |
|  | | Low | | | 0 (0) | 0 (0) | 0 (0) | 0 (0) |
|  | | Moderate | | | 4 (21.1) | 6 (37.5) | 9 (6.9) | 2 (6.9) |
|  | | High | | | 3 (15.8) | 8 (50.0) | 33 (67.4) | 22 (75.9) |
| Metformin, n (%) | |  | | | 2 (10.5) | 3 (18.8) | 23 (46.9) | 9 (31.0) |
| Abbreviations: CKD: chronic kidney disease (stage ≥3; eGFR <60 mL/min/1.73 m²) .; CLTI: chronic limb threatening ischemia; SD: standard deviation. *Low-intensity statin use defined as simvastatin 10 mg or pravastatin 10-20 mg; moderate-intensity as atorvastatin 10-20 mg, rosuvastatin 5-10 mg, simvastatin 20-40 mg or pravastatin 40-80 mg; high-intensity as atorvastatin 40-80mg, rosuvastatin 20-40mg | | | | | | | | |
